# A Fluorogenic-Based Assay to Measure Chaperone-Mediated Autophagic Activity in Cells and Tissues

**DOI:** 10.1101/2023.12.14.571785

**Authors:** Anila Jonnavithula, Megan Tandar, Mohammed Umar, Scott N. Orton, MacKenzie C Woodrum, Sohom Mookherjee, Sihem Boudina, J. David Symons, Rajeshwary Ghosh

## Abstract

**Objective:** Pathologies including cardiovascular diseases, cancer, and neurological disorders are caused by the accumulation of misfolded / damaged proteins. Intracellular protein degradation mechanisms play a critical role in the clearance of these disease-causing proteins. Chaperone mediated autophagy (CMA) is a protein degradation pathway that employs chaperones to bind proteins, bearing a unique KFERQ-like motif, for delivery to a CMA-specific Lysosome Associated Membrane Protein 2a (LAMP2a) receptor for lysosomal degradation. To date, steady-state CMA function has been assessed by measuring LAMP2A protein expression. However, this does not provide information regarding CMA degradation activity. To fill this dearth of tools / assays to measure CMA activity, we generated a CMA-specific fluorogenic substrate assay.

**Methods:** A KFERQ-AMC [Lys-Phe-Asp-Arg-Gln-AMC(7-amino-4-methylcou-marin)] fluorogenic CMA substrate was synthesized from Solid-Phase Peptide Synthesis. KFERQ-AMC, when cleaved via lysosomal hydrolysis, causes AMC to release and fluoresce (Excitation:355 nm, Emission:460 nm). Using an inhibitor of lysosomal proteases, i.e., E64D [L-trans-Epoxy-succinyl-leucylamido(4-guanidino)butane)], responsible for cleaving CMA substrates, the actual CMA activity was determined. Essentially, *CMA activity = (substrate)_fluorescence_ - (substrate+E64D) _fluorescence_.* To confirm specificity of the KFERQ sequence for CMA, negative control peptides were used.

**Results:** Heart, liver, and kidney lysates containing intact lysosomes were obtained from 4-month-old adult male mice. First, lysates incubated with KFERQ-AMC displayed a time dependent (0-5 hour) increase in AMC fluorescence vs. lysates incubated with negative control peptides. These data validate the specificity of KFERQ for CMA. Of note, liver exhibited the highest CMA (6-fold; p<0.05) > kidney (2.4-fold) > heart (0.4-fold) at 5-hours. Second, E64D prevented KFERQ-AMC degradation, substantiating that KFERQ-AMC is degraded via lysosomes. Third, cleavage of KFERQ-AMC and resulting AMC fluorescence was inhibited in Human embryonic kidney (HEK) cells and H9c2 cardiac cells transfected with *Lamp2a* vs. control siRNA. Further, enhancing CMA using *Lamp2a* adenovirus upregulated KFERQ degradation. These data suggest that LAMP2A is required for KFERQ degradation. **Conclusion.** We have generated a novel assay for measuring CMA activity in cells and tissues in a variety of experimental contexts.

Graphical Abstract
Proposed mechanism of KFERQ-AMC degradation via CMA

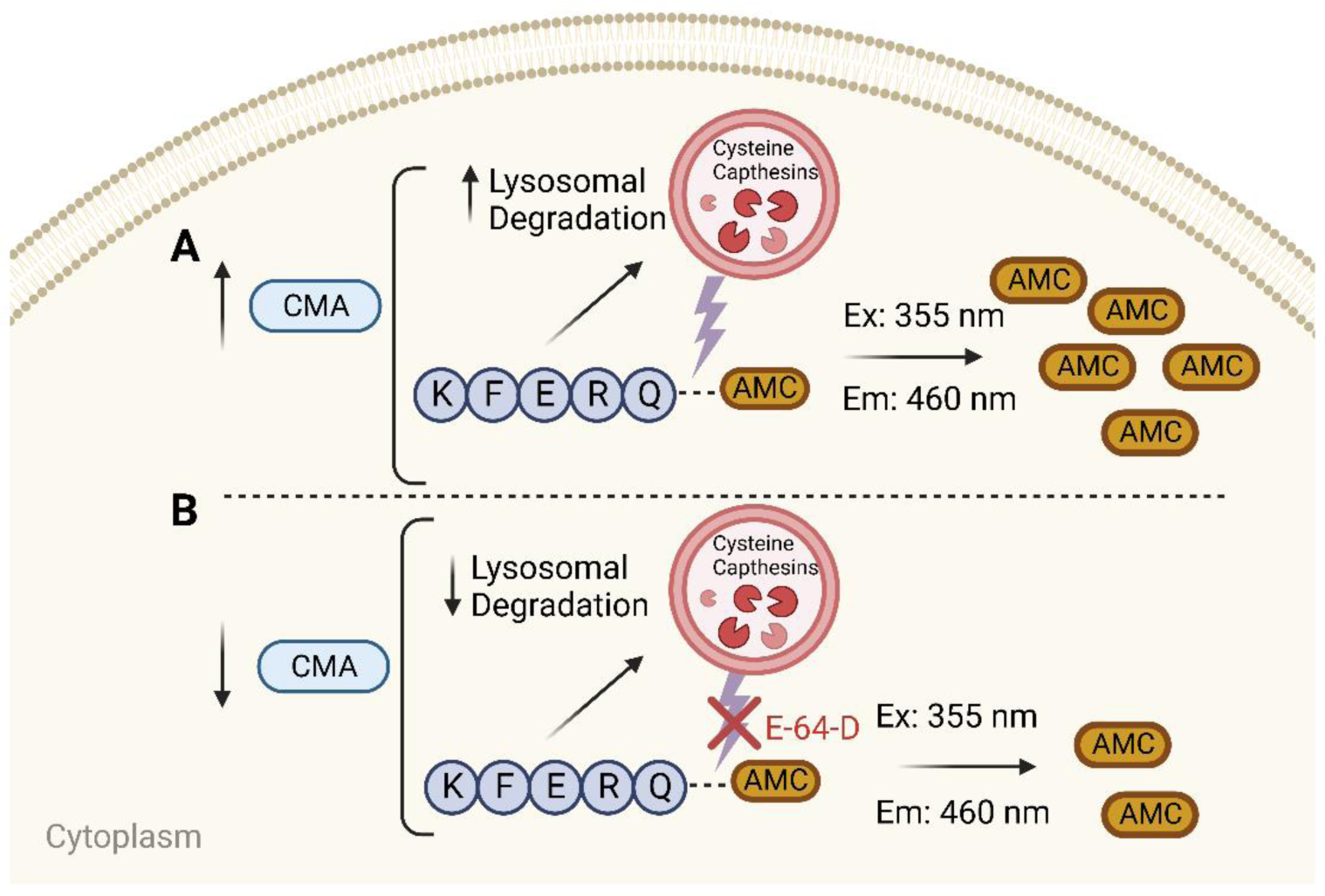

## INTRODUCTION

Tight regulation of protein homeostasis, that is, the balance between protein synthesis and protein degradation, is integral to maintaining normal cellular function and health. Many types of pathologies, including neurodegenerative, cardiovascular, cancer, liver diseases and aging, are associated with dysfunctional proteolytic pathways which can lead to the accumulation of cytotoxic misfolded proteins (1–6). There are two well-known intracellular proteolytic pathways, i.e. the ubiquitin proteasomal pathway and the macroautophagy-lysosomal pathway. Both these pathways maintain normal protein turnover by contributing to bulk protein degradation. Alternately, a third and relatively new protein degradation pathway is the chaperone-mediated autophagy (CMA), which selectively degrades specific substrate proteins which includes misfolded and mutant proteins.

Recent studies have demonstrated the beneficial role of CMA in various pathological conditions such as neurodegenerative diseases, diabetes, and cancer. What makes CMA distinct from UPS or macroautophagy is its ability to selectively recognize and degrade specific protein substrates bearing a unique KFERQ-like motif. CMA employs chaperones to bind targeted dysfunctional proteins for delivery to a CMA-specific Lysosome Associated Membrane Protein 2A (LAMP2A) receptor for lysosomal degradation. In the lysosomal lumen, the substrate proteins are rapidly degraded (within 10 minutes) by lysosomal proteases namely cysteine proteases (cathepsins). To date, LAMP2A expression have been used to assess static CMA function whose levels have been shown to directly correlate with CMA activity in various cells and tissues (7–9). In the past decade, several methods to monitor CMA activity have been established, including measurement of protein degradation rates using radiolabeled amino acid in the protein substrates, isolation of lysosomal membrane vs. matrices and immunoblotting for the CMA components (LAMP2A and HSC70), immunofluorescence to determine the colocalization of LAMP-2A and HSC70 to identify the CMA-active lysosomes in cultured cells, and monitor the direct translocation of known CMA substrates in purified lysosomes from cells or tissues (10). More recently, a photoconvertible fluorescent CMA reporter was developed which allows monitoring CMA in live cultured cells and animals using microscopy-based assay (11, 12). These methods, although reliable, produce certain limitations. For example, radioactivity-based assays present several inherent disadvantages, including radiation hazard, long counting times and high cost of waste disposal (13, 14). Furthermore, owing to the semi-quantitative nature of microscopy, the fluorescent signal of the reporter may not always reveal the precise status of CMA (15). Additionally, while these methods provide information about the steady-state CMA, insights concerning the more relevant end point, i.e. CMA flux, cannot be attained. Therefore, a complementary and versatile method to monitor CMA activity and flux is needed to overcome the limitations of measuring CMA. In this manuscript we have established a novel fluorescent-based CMA activity assay and validated its utility to measure CMA in cells and tissues.

In this protocol, we demonstrate extraction of intact lysosomes from adult mouse hearts, liver, and kidneys, and in cultured rat cardiomyocytes. We next demonstrate the use of a sensitive KFERQ-AMC (Lys-Phe-Asp-Arg-Gln-AMC) fluorogenic substrate to measure CMA activity over time. The substrate KFERQ-AMC is a small peptide attached to an AMC group (7-amino-4-methylcou-marin) that fluoresces at 460 nm when cleaved via hydrolysis from the peptide and excited at 355 nm. The amount of AMC cleavage directly corresponds to CMA activity in the samples. To determine CMA flux, a parallel set of the same samples were treated with a lysosomal inhibitor, trans-Epoxy-succinyl-leucylamido(4-guanidino)butane, (E64D), which blocked KFERQ-AMC degradation. Further negative-control peptides were used to confirm the specificity of KFERQ for CMA. Using this assay, we conducted a comprehensive analysis of CMA flux in various cells and tissues. In conclusion, this protocol is readily adaptable to measure CMA activity accurately and consistently in various cells and tissues, and can be successfully employed to identify different CMA activators/inhibitors as potential drugs for the prevention or treatment of a variety of pathologies.

## RESULTS

### Adult mouse heart, liver, and kidneys show KFERQ-AMC peptide cleavage compared to negative-control peptides

Heart, liver, and kidneys were isolated from 4-month-old adult male C57BL/6J mice (Jackson Laboratories) (6). The animals were handled according to institutional animal care protocol at the University of Utah. The tissue lysates were prepared as described in the Methods. A general workflow of the assay is shown in **Fig. 1**. For measuring CMA activity, the samples were incubated with KFERQ-AMC peptide, and the release of AMC fluorescence due to hydrolysis of KFERQ-AMC was used a direct measurement of CMA activity. KFERQ-AMC showed a time dependent increase (0-5 hour) in AMC fluorescence levels, indicating increasing CMA activity (**Fig. 2A – 2E**). To validate the specificity of KFERQ for CMA degradation, two negative control peptides were tested in parallel experiments. The negative control peptides CFCRN-AMC (negative control 1) (**Fig. 2A, 2C, 2E**) or ANERN-AMC (negative control 2) (**Suppl Fig. 2A, 2C and 2E**) failed to undergo lysosomal hydrolysis, suggesting that KFERQ-like motif is required for CMA-mediated degradation.

**Figure 1.**
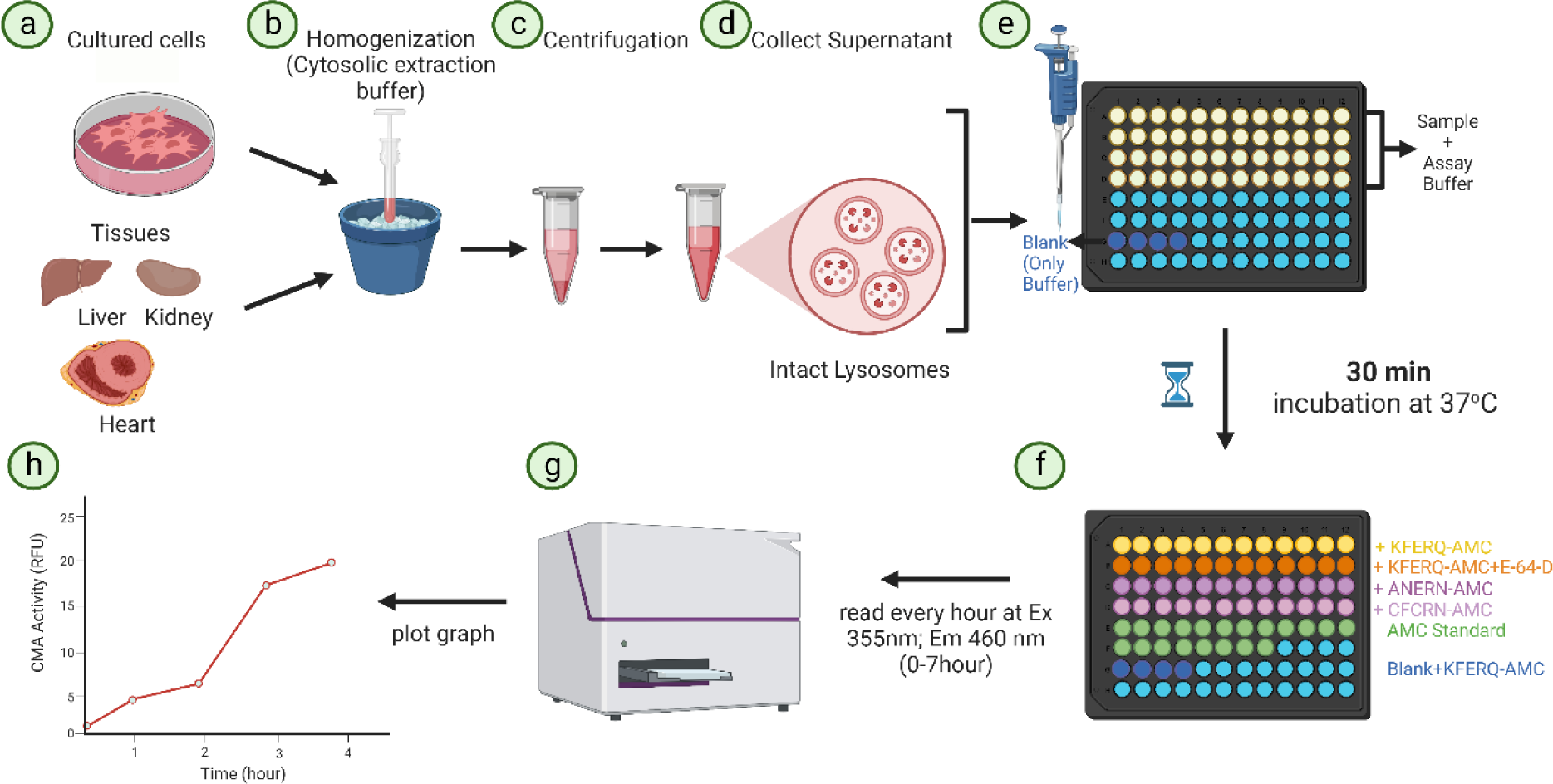
Schematic of the workflow of CMA activity assay

**Figure 2:**
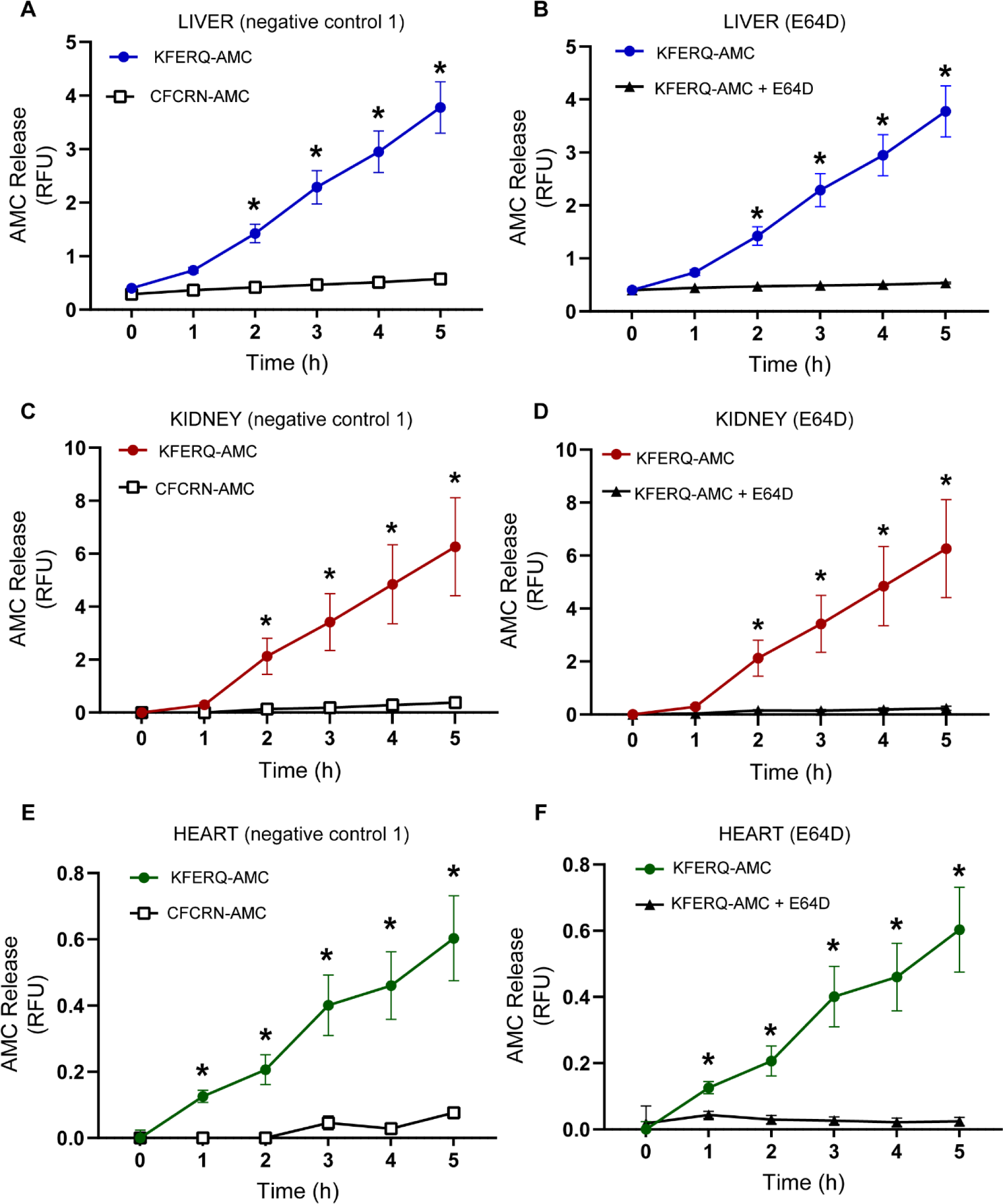
Degradation of KFERQ-AMC vs. CFCRN-AMC (negative control 1) and a lysosomal inhibitor, E64D, in adult mouse heart, liver and kidney. CMA activity indicated by KFERQ-AMC degradation in (A and B, blue line) liver, (C and D, red line) kidney and (E and F, green line) heart. Samples incubated with negative control 1 CFCRN-AMC (A, C, and E, open squares), and the lysosomal inhibitor, trans-Epoxy-succinyl-leucylamido(4-guanidino)butane, (E64D), (B, D, and F, solid triangles) showed no AMC release. For liver n=6, kidney, n=5 and heart n=6. Data are expressed as Mean ± SD. Statistical significance was determined using One Way ANOVA. *p=<0.05 represents the difference between the KFERQ-AMC and CFCRN-AMC or KFERQ-AMC and E64D.

We next determined the efficiency of using a lysosomal inhibitor, E64D, to measure CMA-flux. It is known that lysosomal proteases, namely cysteine proteases, contribute to protein degradation within the lysosomes. E64D is an irreversible, cell permeable inhibitor of lysosomal cysteine cathepsin proteases. It should be noted that all cathepsins (cathepsin B, C, F, K, L, O, S, and X) can be inhibited by E64D (16). We found that addition of E64D successfully inhibited the cleavage of KFERQ-AMC, suggesting that CMA’s degradation capacity was impaired due to the E64D mediated inhibition of lysosomal proteases (**Fig. 2B, 2D, 2F**). Furthermore, since both CMA and macroautophagy utilize lysosomes for protein/organelle degradation, we determined if E64D alters LAMP2A or the macroautophagy marker, LC3II, protein levels. Treatment of H9c2 cardiac cells with E64D did not change either LAMP2A or LC3II levels (**Suppl Fig. 3A-3C**). This data validated that E64D does not inhibit macroautophagy or have any effect on the membrane LAMP2A protein and thus can be utilized to determine CMA flux in tissues and cells.

**Figure 3.**
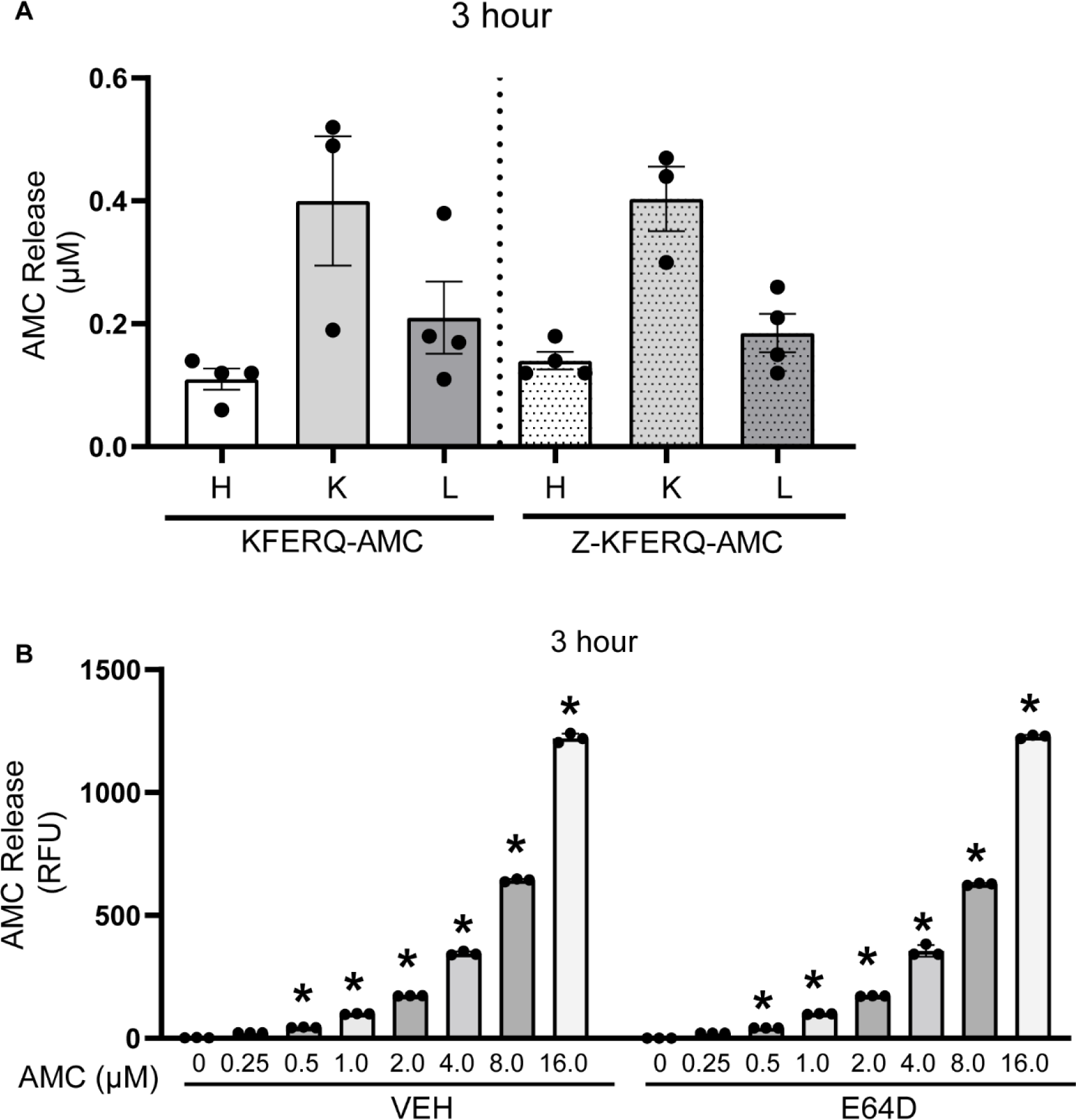
KFERQ-AMC emits no background fluorescence and E64D does not directly influence AMC fluorescence. (A) Heart liver and kidney were incubated with 50 μM of either KFERQ-AMC or Z-KFERQ-AMC for 3 hours. The bar graph represent CMA activity at 3 hours. n = 4/group. (B) Different concentrations of AMC (0-16 μM) were incubated with 5 mM E64D or VEH (DMSO) for 3 hours. The bar graph represents the amount of AMC fluorescence in the presence or absence of E64D. Data are expressed as Mean ± SD. Statistical significance obtained using one-way ANOVA for (A) and unpaired t test for (B). For (A), *p=<0.05 represents the difference between the KFERQ-AMC and Z-KFERQ-AMC; for (B), *p=<0.05 represents the difference between the VEH or E64D groups between no AMC (0μM) and the AMC groups.

We included blank wells as additional negative controls to evaluate if any background KFERQ-AMC fluorescence is generated. No substrate reaction was observed in the blank wells (**Suppl Fig. 2B, 2D and 2F**), suggesting that the AMC cleavage was due to the occurrence of CMA-mediated degradation of KFERQ-AMC. CMA activity of each sample was determined by comparing the fluorescence of cleaved AMC to the known concentration of free AMC using an AMC standard curve (an example of AMC standard curve is shown in **Suppl Fig. 1E**). These data demonstrate the efficacy of employing KFERQ-AMC as a bonafide CMA substrate to measure CMA activity and function in different types of tissues.

### KFERQ-AMC emits no background fluorescence

N-terminal Z (carbobenzyloxy) and C-terminal AMC are sometimes paired so that the peptide has a low background fluorescence until it is hydrolyzed releasing free AMC, which fluoresces. It is also known that peptides without the N-terminal Z works well because of the drastic increase in fluorescence when AMC is released. Nevertheless, for each experiment, we have validated the KFERQ-AMC fluorescence before (0 hour) and in the blank which showed minimum or no fluorescence vs. after protease hydrolysis of the peptide (**Suppl Fig 2B, 2D and 2F**). Furthermore, we generated a Z-KFERQ-AMC peptide and compared its fluorescence levels to KFERQ-AMC in heart, liver, and kidney from adult mice (Fig 3A). The levels of AMC fluorescence in Z-KFERQ-AMC were similar to KFERQ-AMC in the tissues tested at 3 hours (Fig 3A).

### E64D does not directly inhibit AMC fluorescence

To determine if E64D directly binds to AMC and interferes with its fluorescence levels, we incubated different amounts of AMC (0-16 μM) with or without E64D and measured AMC fluorescence for 3 hours. At 3 hours, the amount of AMC fluorescence in the presence of E64D was similar to AMC not incubated with the lysosomal inhibitor (**Fig. 3B**). These results indicate that E64D does not directly influence AMC fluorescence and rather acts by inhibiting lysosomal function to impede KFERQ-AMC degradation.

### CMA activity correlates to LAMP2A protein levels

If the CMA-specific receptor, LAMP2A, is both necessary and sufficient for CMA function as shown before (7, 9, 17), LAMP2A protein levels should directly correlate with KFERQ-AMC degradation (10, 17). Equal amounts (∼35 μg) of liver, heart and kidney proteins were separated by gel electrophoresis and immunoblotting was performed as described previously (7). Compared to the heart, LAMP2A level was higher in kidney (∼8-fold) followed by liver (∼6-fold) (**Fig. 4A and 4B**). Consistently, CMA **flux** determined by the amount of KFERQ-AMC cleavage in the presence and absence of E64D was found to be maximum in the kidney, followed by liver and heart (**Fig. 4C**). These data suggest lysosomal degradation of KFERQ-AMC is regulated by LAMP2A levels.

**Figure 4.**
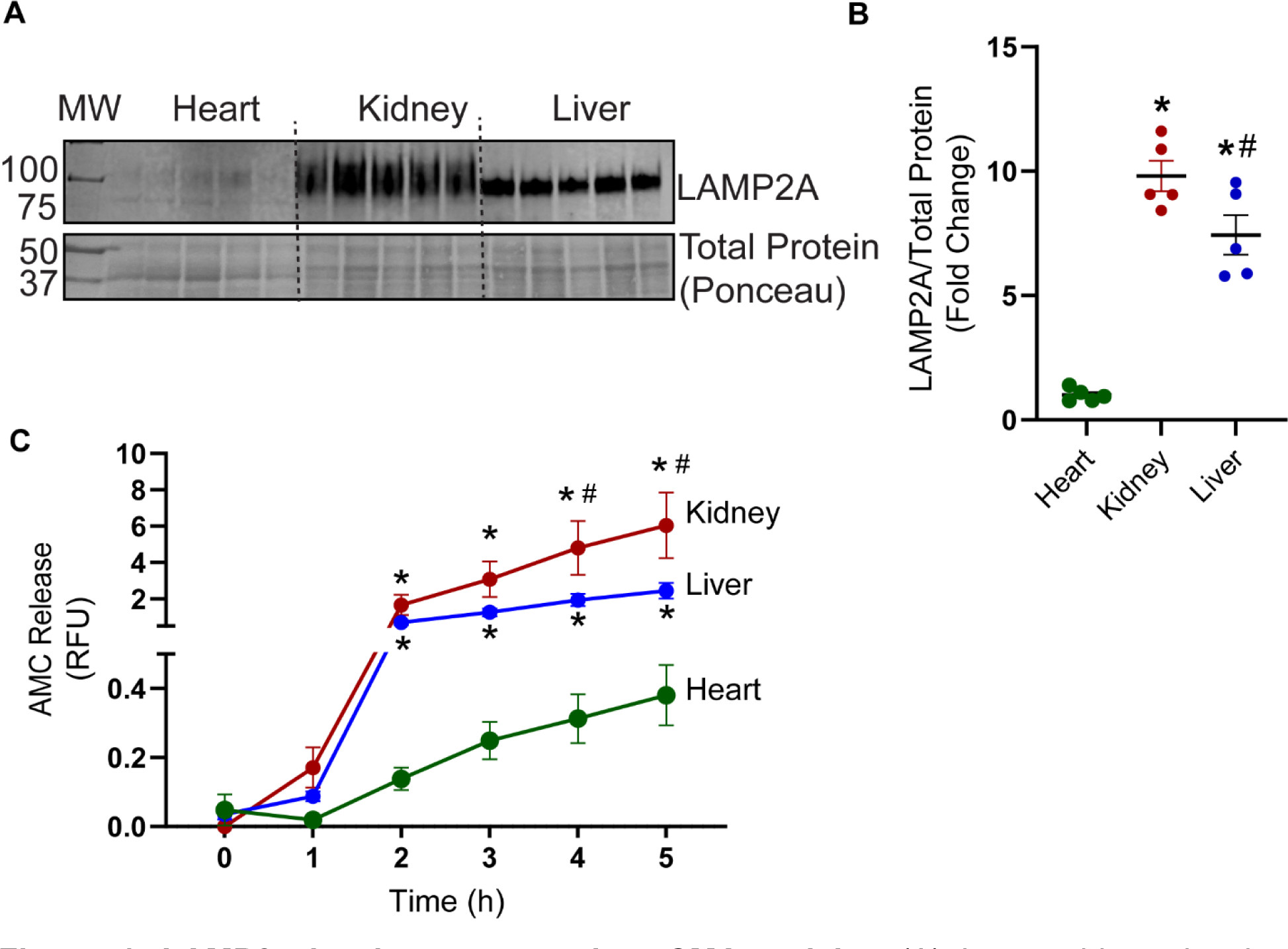
LAMP2a levels correspond to CMA activity. (A) Immunoblots showing LAMP2A protein levels in the kidney liver and heart and (B) densitometry analysis of LAMP2A normalized to total protein levels. (C) KFERQ-AMC degradation in the kidney, liver, and heart. n=5 for all groups. The corrected values after blank subtraction have been deducted from E64D group to determine CMA flux. Data are expressed as Mean ± SD. Statistical significance between the groups was determined by one-way ANOVA. *(p<0.05) represents the difference between heart and liver or kidney; ^#^(p<0.05) represent the difference between kidney and liver.

### CMA flux declines with *Lamp2a* silencing in cultured cells

To further validate that the cleavage of KFERQ-AMC occurs via CMA, we performed *Lamp2a* silencing in cultured HEK 293 and H9c2 rat cardiomyoblasts (7). Immunoblotting validated successful knockdown of LAMP2A (∼90%) in both HEK 293 and H9c2 cells vs. the control siRNA transfected cells (**Fig. 5A-B and Suppl Fig. 4A-B**). Further, silencing of LAMP2A reduced KFERQ-AMC degradation as observed by decreased AMC fluorescence (**Fig. 5C** and **Suppl Fig. 4C**). These results confirm that LAMP2A is required for the uptake of KFERQ-AMC into the lysosomes for its subsequent degradation via CMA.

**Figure 5.**
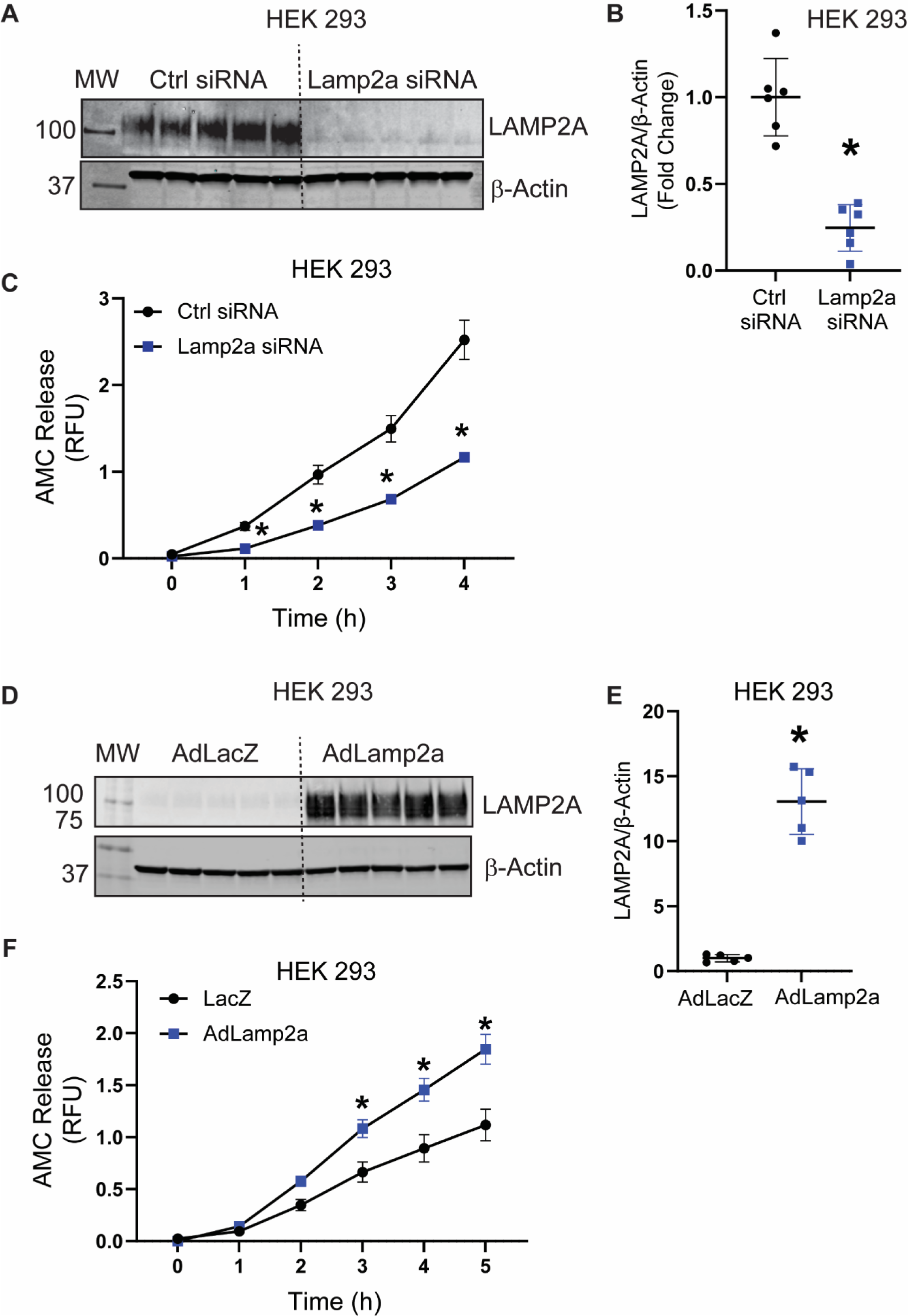
LAMP2A suppression inhibits, and overexpression enhances CMA activity. *Lamp2a* gene was silenced in human embryonic kidney 293 (HEK 293) cells using a *Lamp2a* siRNA. Control (Ctrl) groups were transfected with a negative control siRNA. Alternatively*, Lamp2a* gene was overexpressed in HEK 293 cells using a *Lamp2a* adenovirus (*AdLamp2a*) encoding the *Lamp2a* gene. A LacZ group (*AdLacZ*) was added as a negative control group. Immunoblots show LAMP2A protein in HEK 293 (A and D). (B) and (E) are the respective densitometries of LAMP2A normalized to β-Actin levels. KFERQ-AMC degradation was measured in HEK 293 (C and F). The corrected values after blank subtraction have been deducted from E64D group to determine CMA flux. n=6 for HEK 293. Data are expressed as Mean ± SD. Statistical significance between the groups was determined by one-way ANOVA. *(p<0.05) represents the difference between the Ctrl sirna vs, *Lamp2a* sirna or Ad*LacZ* vs. Ad*Lamp2a* groups.

**Figure 6.**
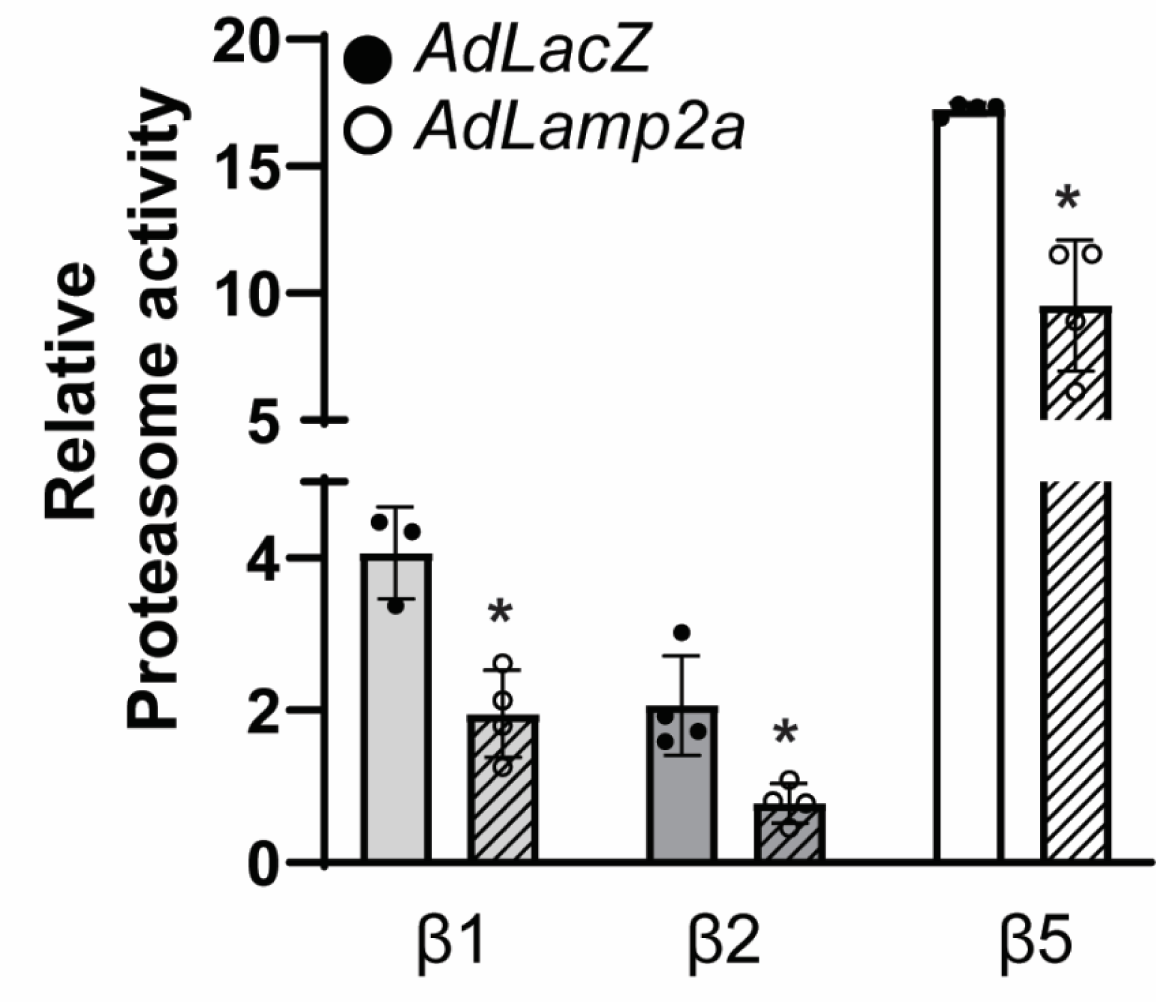
LAMP2A overexpression decreases proteasome activity. *Lamp2a* gene was overexpressed in human embryonic kidney 293 (HEK 293) cells using a *Lamp2a* adenovirus (*AdLamp2a*) encoding the *Lamp2a* gene. A LacZ group (*AdLacZ*) was added as a negative control group. Proteasomal activity (β1, β2 and β5) were measured in the cells. n=4 for HEK 293. Data are expressed as Mean ± SD. Statistical significance between the groups was determined by t-test between the two groups for each activity. *(p<0.05) represents the difference between *AdLamp2a* and *AdLacZ* groups.

### CMA flux increases with *Lamp2a* overexpression in cultured cells

We next confirmed if upregulation of LAMP2A increases CMA-mediated KFERQ-AMC cleavage. LAMP2A protein was upregulated by infecting HEK 293 cells with adenovirus encoding the *Lamp2a* gene (7). Immunoblots showed a successful increase in LAMP2A levels in HEK 293 cells (**Fig. 5D-E**). Consistently, LAMP2A upregulation correlated with an increase in AMC fluorescence HEK 293 cells (**Fig. 5F**), indicating increased KFERQ-AMC degradation and consequently enhanced CMA function. These results confirm that LAMP2A levels regulate the degradation of CMA substrates.

### Increased CMA activity is associated with decreased proteasome function

In a recent study we demonstrated that enhancing CMA using *Lamp2a* adenoviruses increased macroautophagic activity in primary neonatal rat cardiomyocytes (7). Herein, we determined the effect of *Lamp2a* overexpression on proteasome function in HEK cells. Using substrates specific for the three proteasomal catalytic sites required for the initiation of the reaction, chymotrypsin (β5), trypsin (β2) and caspase (β1)-like activities were determined in HEK cells overexpressing *Lamp2a* (18). As expected, β5 activity (the most active catalytic site of the proteasome) was notably higher relative to β1 (4-fold) or β2 (8-fold) activities in the *LacZ* control cells. Enhancing CMA using *Lamp2a* adenoviruses, attenuated all the three proteasomal activities (β5:1.8-fold; β1:2-fold; and β2:2.8-fold). These results suggest that a cross talk exists between CMA and proteasome function where inducing the CMA activity impairs proteasome function which is demonstrated by the loss of all three catalytic subunit proteasomal activities.

## MATERIALS AND METHODS

The CMA activity assay would take approximately 9-10 hours including sample preparation and running the plates.

1. Dounce Homogenizer
2. Temperature controlled microcentrifuge
3. 96 well black, flat-bottom plates.
4. Temperature controlled microplate reader capable of reading fluorescence. For our experiments, we used SkanIt Software 6.1.1 RE, ver. 6.1.1.
5. <u>CMA Extraction Buffer (CEB)</u>

a. 50 mM Tris-HCl, pH 7.4 (Sigma Aldrich, Cat. # T2319)
b. 0.5 mM Ethylenediaminetetraacetic acid (EDTA) (Sigma Aldrich, Cat. #E5134)
c. 2 mM Adenosine Triphosphate (ATP) (Sigma Aldrich Cat. # A1852)
d. 1 mM Dithiothreitol (DTT) (Roche Cat. # 10197777001)
e. 250 mM Sucrose (Sigma Aldrich, Cat. # S0389)
f. 5 mM Magnesium Chloride (MgCl_2_) (Sigma Aldrich, Cat no.# M8266)
g. 0.25% Digitonin (Sigma Aldrich Cat. # D141)
h. Nanopure Water

*\*\* Protein isolation was done using non-denaturing buffer conditions to maintain functional CMA. We used Tris, DTT and glycerol containing buffer in the presence of a mild detergent (Digitonin) to disrupt the cell membrane. This lysis buffer was tested to generate favorable outcome. The extraction buffer should be made fresh. Higher stock concentration of the buffers can be prepared and stored at room temperature (Tris-HCl, EDTA, MgCl_2_) or at −20 °C (ATP, Sucrose, Digitonin). On the day of the experiment, the buffers can be diluted to the desired concentrations. DTT must be prepared fresh*.

6. CMA Assay Buffer (CAB)

a. 50 mM Tris-HCl, pH 7.4
b. 40 mM Potassium Chloride (KCl)
c. 5 mM MgCl_2_
d. 0.5 mM ATP
e. 1 mM DTT
f. 0.05mg/ml Soybean Trypsin Inhibitor (ThermoFisher Cat. # 17075029)
g. Heat shock 70 kDa protein 8 Lipopolysaccharide-associated protein 1 (HSC 70, Abcam Cat. # AB78431)
h. Nanopure Water
7. <u>CMA substrates</u>: KFERQ-AMC and Scrambled peptides (ANERN-AMC and CFCRN-AMC) (LifeTein, LLC. Hillsborough, NJ): Prepare a 10 mM stock of each substrate in DMSO. Aliquot and store the stocks at −20 °C. The working concentration for each substrate tested was 50 μM. *<u>Note 1</u>. The negative-control peptides were designed using Combinatorial 3-Positional Scan, Mimotopes, The Peptide Company. (*http://www.mimotopes.com/peptideLibraryScreening.asp?id=93&e=captcha).
8. E64D (Enzo Life Sciences, BML-PI107): Prepare a stock of 10 mM stock in DMSO. Aliquot and store at −20 °C. The working concentration should be 5 mM.
9. AMC (7-amino-4-methylcoumarin) fluorescence reference standard (ThermoFisher Scientific, A191): Prepare a 100 mM stock in DMSO. Prepare aliquots of the stock and store at −20 °C.

## 3. METHODS

A schematic of the workflow is shown in Figure 1.

### 3.1 Tissue Homogenization (*Estimated Time: 5 minutes/sample*)

*All procedures should be carried out on ice, unless specified otherwise*.

3.1.1 Frozen tissue samples can be used for measuring CMA activity. Weigh approximately 25 mg of each tissue sample and pulverize using a mortar and pestle while keeping the samples chilled in liquid nitrogen.
3.1.2 Transfer the pulverized samples to pre-chilled 1.5 ml Eppendorf tubes and add 500 μl CEB.
3.1.3 Homogenize the tissue samples using a Dounce homogenizer on ice.
3.1.4 Approximately, 25 strokes should be applied to disrupt the tissues.
3.1.5 Centrifuge the samples at 12,000 x g for 10 minutes at 4°C. Next, transfer the supernatant to a clean pre-chilled 1.5 ml tube. The extracted samples can be used directly for the assay or stored at −20 °C.

*\*\* Centrifugation step may be repeated if the solution appears cloudy*.

*Do not snap freeze or vortex the samples. The samples should be used within 1 week of extraction*.

### 3.2 Cell Homogenization (Estimated Time, 5 minutes/sample)

3.2.1 Cells were maintained in 10 cm dishes. On the day of experiment, transfer the plates to ice. Wash the cells with pre-chilled 1X PBS, twice. Add 500 μl of CEB to each plate and incubate for 5 minutes on ice.
3.2.2 Using cell scrapers, scrape the cells and transfer to a Dounce homogenizer.
3.2.3 apply 20 strokes to disrupt the cell membrane.
3.2.4 Centrifuge the samples at 12,000 x g for 10 minutes at 4°C. Carefully transfer the supernatant into clean pre-chilled 1.5 ml tubes. The extracted samples can be used directly for the assay or stored at −20 °C.

### 3.3 Protein Determination and Immunoblotting (Estimated Timing: 2 days)

<u>4</u> Quantify the protein concentrations of the CEB extracted lysates using a protein determination assay. For our studies, we performed Bradford assay (BioRad, 5000006) for protein quantification against a bovine serum albumin standard curve (My BioSource, MBS355528). The samples were scaled to equal concentrations (1 μg/μl), and 20-40 μg samples were used for running the assay.

### CMA Activity Assay

#### Running the assay plate (5-7 hours)

Our results indicate that sensitive and reproducible CMA activity measurements can be obtained with 40 μg. However, a pilot experiment is recommended using different amounts (10, 20, 30, and 40 μg) of tissue or cell lysates in a 96-well black well.

- Using a repeater pipette, add 2 ul of inhibitor (E-64-D) to the appropriate well. For the control wells add 2 ul DMSO.
- Add CAB to each well (200 μl - V _lysate_ - V _substrate_ - V _inhibitor_); (V=Volume).
- Add 20 μl (40 μg) lysate to each well. Include 3-4 blank wells [(buffer + substrate)-sample]. *Note: For multiple plates, stagger addition of inhibitors by 15 minutes*.
- Seal the plates with a clear film, shake the plate briefly (5 seconds) on a flat surface to avoid spill, and incubate for 30 minutes at 37°C.
- Take a fluorescent reading immediately at 355 nm excitation and 460 emission.

Continue reading the plate with 1-hour increments for 5-7 hours.

*Note: Set the plate reader to be preheated at 37°C before taking readings. If more than one plate is run, stagger the measurements to have identical time of measurement since the addition of substrate*.

#### AMC Calibration Curve

- An AMC standard curve should be simultaneously prepared to determine the CMA activity of the unknown samples by comparing the concentrations of the unknown samples to the known concentration of AMC.

1. Thaw a vial containing 100 mM free AMC stock and dilute to make a 50 μM AMC stock solution in CAB buffer.
2. Make the following serial dilutions of AMC to final concentrations of: 16, 8,4, 2, 1, 0.5. 0.25 and 0 μM AMC in the same black well plate (Supplemental Figure 1E).
3. Plot a standard curve of AMC fluorescence and determine the unknown concentration of free AMC released due to KFERQ-AMC cleavage.

### CMA Activity Calculation

CMA activity is represented as Relative fluorescence units (RFU). The relative CMA activity is calculated based on the amount of free AMC in relative fluorescence unit (RFU)] released from KFERQ-AMC by lysosomal cleavage, determined by comparing to the standard calibration curve of AMC (Supp Fig 1). AMC standard curve should be prepared during each experiment. The mean RFU value for each sample in duplicates should then be calculated. The mean values of all samples should be corrected by subtracting the value of the blank. The actual CMA flux can be measured by subtracting each sample from the E64D treated sample.

### Additional Established Methods

#### H9c2 and HEK 293 cell culture

H9c2 immortalized cardiomyoblasts isolated from the embryonic rat heart tissue (ATCC; Passage no. 4 to 10) and human embryonic kidney 293 (HEK 293) cells (ATCC; Passage no. 4 to 25) were utilized to determine KFERQ-AMC degradation. The cells were maintained in 75 ml cell culture flasks using high glucose Dulbecco’s modified Eagle’s medium (DMEM) high glucose, pyruvate, (ThermoFisher Scientific, 11995-065) at 37°C and 5% CO_2_, with 10% fetal bovine serum (FBS), supplemented with and 1% penicillin/streptomycin (complete growth media).

Once the cells reached a confluency of 70-80%, they were used for various experiments.

#### E64D treatment

H9c2 cells were treated with 5 mM E64D for 4 hours. The cells were then collected for performing immunoblots.

#### *Lamp2a* silencing and overexpression

Construction of a *Lamp2a* siRNA has been described before (7). H9c2 or HEK293 cells were transfected with 12 nM of a *Lamp2a* siRNA (Ambion Life Technologies) or a medium GC control siRNA (Thermo Fisher, Cat. No. 4390843) was used as a negative control using Lipofectamine RNAi (4 µg/µl) (ThermoFisher Scientific, Cat. No. 13778150) in Opti-MEM media (ThermoFisher Scientific, Cat. No. 31985070) for 24 hours. Next, siRNA containing media was replaced with DMEM growth media supplemented with 10% FBS and 1% penicillin/streptomycin. After 48 hours, the cells were collected for performing CMA activity assay and immunoblotting.

HEK293 cells were infected with adenoviral constructs expressing either *Lamp2a*, to induce CMA activity, or a negative-control LacZ, as described before (7). Briefly, HEK293 cells were infected with 120 MOI of adenovirus in serum deficient DMEM for 24 hours. The adenovirus-containing media was replaced with DMEM, supplemented with 10% FBS, and 1% penicillin/streptomycin. 48 hours later, the cells were collected for downstream applications.

#### Immunoblotting for LAMP2A protein

In addition to running the CMA activity assay, the samples were separated by SDS-PAGE and immunoblotted to determine LAMP2A protein levels as described previously (7). The CEB lysates were mixed with Laemmli SDS sample buffer (Alfa Aesar, J61337-AC). The samples were next boiled at 95°C for 5 minutes and 40-60 µg of each protein sample was separated using 4-15% TGX gels (BioRad, 5671084). A molecular weight marker was also loaded into one lane simultaneously for each gel (LICOR, 928-40000). For gel electrophoresis, 1X Tris/Glycine/SDS buffer (BioRad, 1610732) running buffer was used and the gel was run at 100 volts for ∼1.5 hours. The proteins were next transferred to 0.2 µm PVDF membranes (BioRad, 1620239) using Biorad Trans-Blot Turbo Transfer System (2.5 Ampere, 25 Volt, 7 minutes). Membranes were blocked using Aquatic Block-azide free PBS (Arlington Scientific, Inc.) for 1 hour at room temperature, and probed overnight at 4⁰C with LAMP2A (1:200; AMC2; Invitrogen, 512200), or β-Actin (1:1000; AC-15; Sigma-Aldrich, A5441) as the loading control. For samples showing variations in β-Actin, total protein stained with Ponceau stain was used as loading control. The next day, the membranes were washed in PBS-T containing 0.01% Tween and incubated with species–specific secondary antibodies conjugated to infrared dyes (1:10,000; LICOR, 926-3221; 926-68070) for 1 hour at room temperature. The blots were imaged in a LICOR Odyssey CLx scanner and the quantification of the bands was done using the ImageStudio 2.0.38 software.

#### Proteasome activity measurement

Proteasome assays were carried out as described before (18, 19). Briefly, the assay was performed in the absence or presence of a specific proteasome inhibitor: bortezomib. Cultured cells were homogenized and centrifuged at 12,000 x g for 15 mins at 4°C. Supernatant containing the proteasomes were extracted and protein concentration determined. 20 µg of the samples were used for measuring the catalytic activities: β1, β2 and β5. The reaction for each assay was initiated using the specific substrates (Enzo Life Sciences, NY, USA): β1 substrate Z-LLE-AMC; β2 substrate Boc-LSTR-AMC and β5 substrate Suc-LLVY-AMC.

#### Statistical Analysis

The data are presented as mean ± standard deviation of the mean. All statistical analyses were performed using GraphPad Prism software version 9. The value of p < 0.05 was considered statistically significant for all the experiments. Significant differences between two groups were evaluated using an unpaired two-tailed t-test. A one-way ANOVA was performed to determine differences in mean values across the different time points between two groups. Tukey’s multiple comparisons post hoc test was used to identify the differences between the two experimental groups. Information pertaining to the statistical tests is included in each legend.

## DISCUSSION

We have generated a novel fluorogenic assay that can successfully measure CMA activity in a host of cells and tissues. A novel fluorogenic substrate was designed by linking KFERQ to AMC and lysosomal-mediated degradation of the substrate measured. Enhanced CMA would result in an increase in KFERQ-AMC cleavage causing the free AMC to fluoresce. Alternately, suppression of CMA would lead to reduced KFERQ-AMC degradation resulting in minimum AMC fluorescence. Similar to measuring macroautophagy flux using the macroautophagy inhibitors, *Bafilomycin A1* or chloroquine, we have employed a lysosomal inhibitor, E64D, to measure CMA flux in the samples. Addition of E64D completely blocked KFERQ-AMC degradation due to the inhibition of lysosomal protease activity. E64D did not seem to influence the macroautophagy marker LC3II. This may be because, it functions by suppressing cysteine proteases within the lysosome and blocks lysosomal degradation without affecting LC3II synthesis or macroautophagy. Thus E64D could be used to specifically inhibit CMA to determine CMA flux in cells and tissues. To further validate that KFERQ-AMC peptide could be used as an indicator of CMA activity, we employed genetic models to test that degradation of KFERQ-CMA is CMA-dependent. Suppressing CMA by silencing *Lamp2a* inhibited KFERQ-AMC degradation indicative of decreased CMA activity, while upregulating CMA via *Lamp2a* overexpression, increased AMC fluorescence due to enhanced KFERQ-AMC cleavage which denoted an increase in CMA activity. Using this assay, we were also able to compare the relative proteasomal activities in the *Lamp2a* overexpressing cells. While KFERQ-AMC degradation was high in the *Lamp2a* transduced cells, the catalytic activities of proteasome was significantly impaired, suggesting that at a given time, boosting CMA pathway alone is sufficient to maintain cellular homeostasis and function.

LAMP2A protein levels are known to be both necessary and sufficient for CMA function (7, 9, 10). While this is true, the amount of LAMP2A proteins may not always accurately reflect the actual CMA activity. Our data showed that while *Lamp2a* overexpression caused more than 20-fold increase in LAMP2A protein level, CMA activity was increased by ∼2 fold. It has been shown that activated lysosomal p38 MAPK phosphorylates LAMP2A, causing its accumulation and oligomerization on the lysosomal membrane which is required for substrate translocation to the nucleus (20). We believe that at a given time only a subset of the LAMP2A population assume an active oligomeric conformation and can facilitate substrate translocation to the lysosome for degradation. Thus, the amount of KFERQ-AMC cleavage denotes the actual CMA activity, which may not always be equivalent to the total LAMP2A protein levels.

While there is an array of methods available to measure CMA activity in cells and tissues, most of them rely on expensive live cells/animal models to perform the studies (11). These methods may not be useful to measure CMA in frozen cells and tissues (especially those derived from human), which otherwise may provide valuable information about the status of this specific protein quality pathway. We herein introduce a cost-effective and novel instance that overcomes many of the limitations of measuring CMA displayed in other methods. With the ability to accurately and easily measure CMA, our assay will contribute to advancing the field of protein quality control in a variety of pathophysiological conditions.

## ACKNOWLEDGMENTS

This work is supported by American Heart Association Career Development Award (AHA Award Number: 941327) American Heart Association and RTW Charitable Foundational Grant to RG. SB is supported by grants R01HL149870 and R01HL167866-01A1 from the National Heart, Lung and Blood Institute and grant R01DK128819 from the National Institute of Diabetes and Digestive and Kidney Disease, JDS is supported by RO1HL141540 and SM by an American Heart Association Pre-Doctoral award (AHAPRE1025910).

## CONTRIBUTIONS

AJ and MT prepared the buffers and tissue samples. AJ, MT, MU, SO and MW performed the immunoblots. AJ, MT and MU assisted with data analyses. R.G., AJ, MT and SO wrote the manuscript. RG, SB and JDS edited the manuscript. SM helped with running the assay. R.G. conceived the project, performed the experiments, acquired funding and supervised the project. All authors have read and approved the final manuscript.

## Conflict of Interest

A provisional patent on “A Fluorogenic-Based Assay to Measure Chaperone Mediated Autophagy Activity/Flux in Cells and Tissues”, has been filed by the University of Utah Technology Licensing Office (U-7517).

---

**Supplemental Figure 1:**
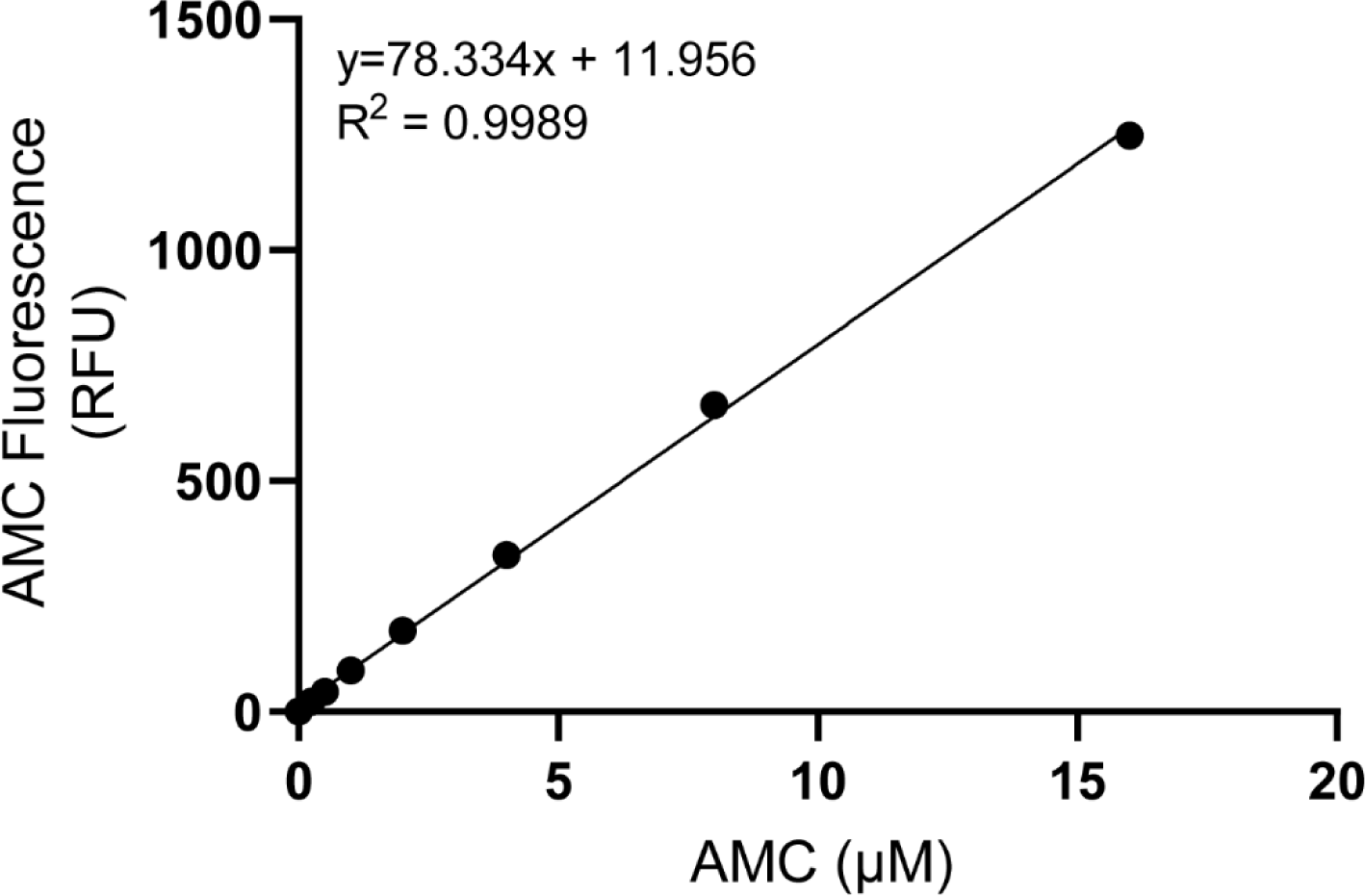
AMC standard curve generation. AMC standard curve was generated using increasing concentrations of AMC (0 μM −16 μM). The amount of AMC fluorescence was used to determine the unknown concentration of AMC release due to KFERQ-AMC cleavage, in the samples.

**Supplemental Figure 2:**
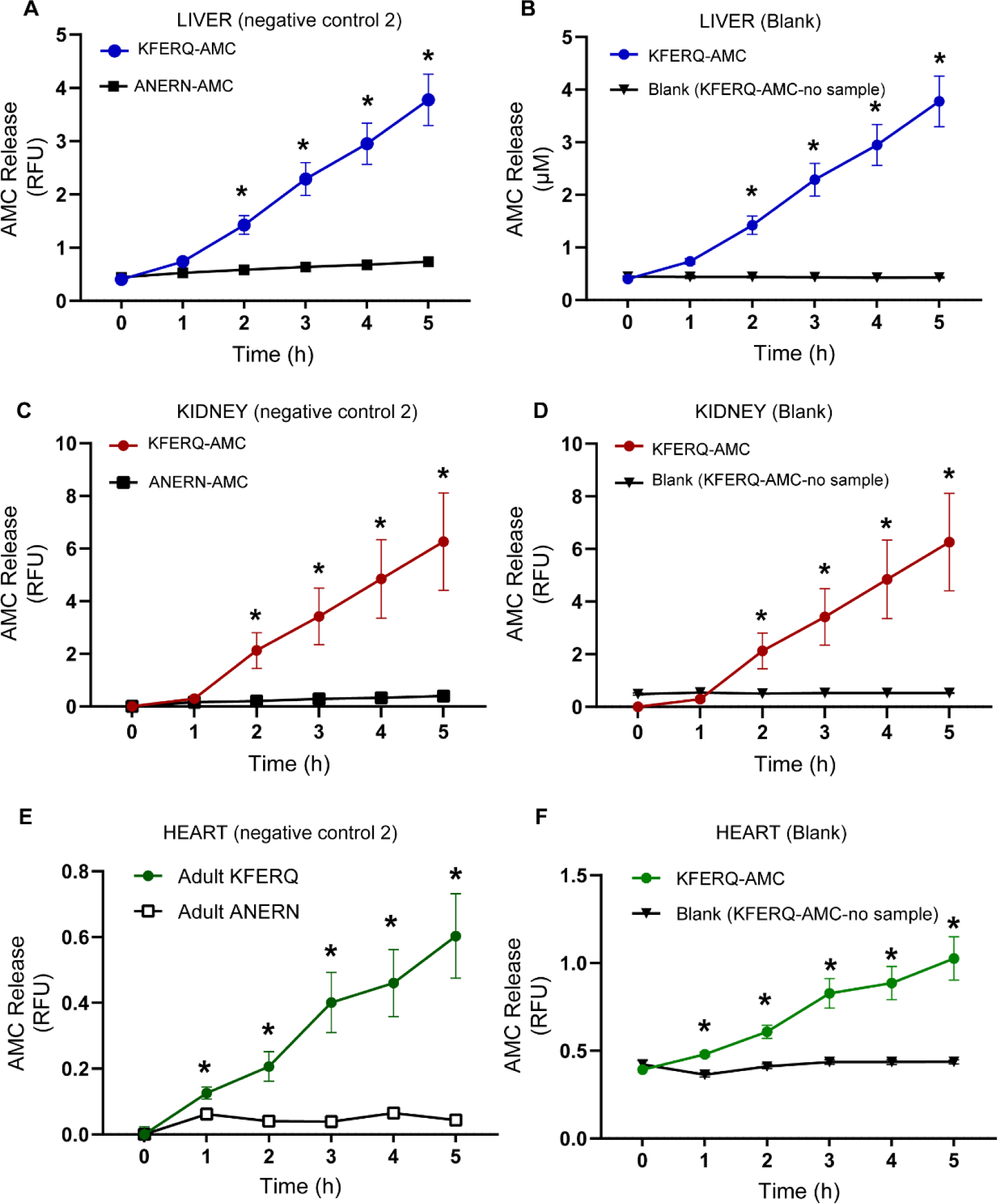
Degradation of KFERQ-AMC vs. ANERN-AMC (negative control 2) and blank in adult mouse heart, liver and kidney. CMA activity indicated by KFERQ-AMC degradation in (A and B, blue line) liver, (C and D, red line) kidney and (E and F, green line) heart. Samples incubated with negative control 1 ANERN-AMC (A, C, and E, solid squares), and blank (buffer only with KFERQ-AMC), (B, D, and F, solid inverted triangles) showed no AMC release. For liver n=6, kidney, n=5 and heart n=6. Data are expressed as Mean ± SD. Statistical significance was determined using One Way ANOVA. *p=<0.05 represents the difference between the KFERQ-AMC and ANERN-AMC or KFERQ-AMC and blank.

**Supplemental Figure 3.**
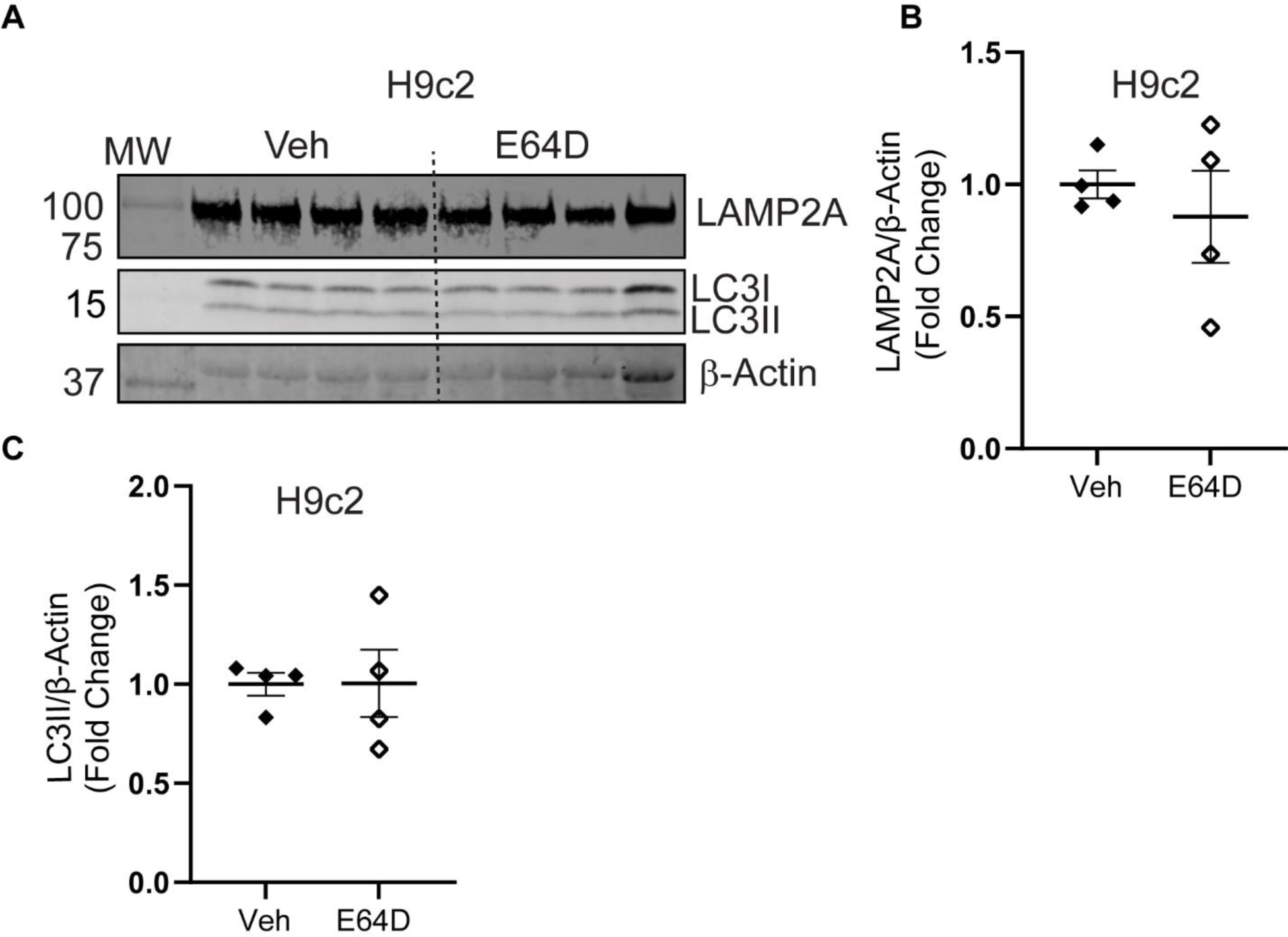
E64D does not affect LAMP2a and LC3II protein levels or AMC fluorescence. H9c2 cells were incubated with 5mM E64D or Vehicle (Veh; DMSO) for 4 hours. (A) Immunoblots show LAMP2A, LC3I and LC3II protein levels. Densitometry of (B) LAMP2a and LC3II (C) normalized to β-Actin levels. n=4/group. Unpaired t-test was performed to determine the difference between the Veh or E64D groups.

**Supplemental Figure 4.**
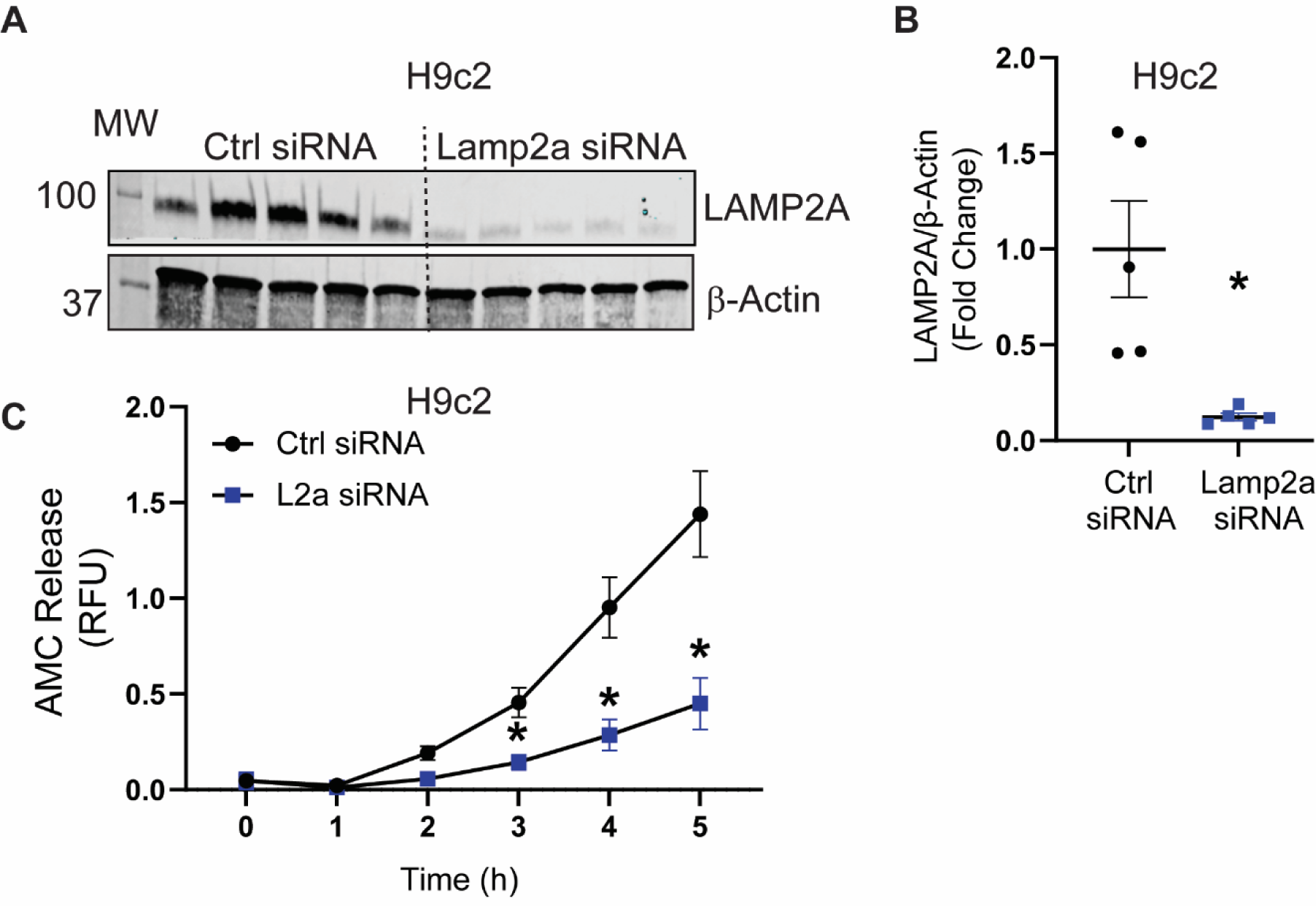
LAMP2A suppression inhibits CMA activity in H9c2 cardiac cells. *Lamp2a* gene was silenced in H9c2 cells. (A) Immunoblots show LAMP2A protein in HEK 293. (B) is the respective densitometry of LAMP2A normalized to β-Actin levels. (C) KFERQ-AMC degradation was measured. The corrected values after blank subtraction have been deducted from E64D group to determine CMA flux. n=6; data are expressed as Mean ± SD. Statistical significance between the groups was determined by one-way ANOVA. *(p<0.05) represents the difference between the Ctrl sirna vs, *Lamp2a* sirna groups.

## Notes

### Competing Interest Statement

A provisional patent has been filed by the University of Utah Technology Licensing Office (U-7517).

